# Estimation of Membrane Curvature for Cryo-Electron Tomography

**DOI:** 10.1101/579060

**Authors:** Maria Kalemanov, Javier F. Collado, Wolfgang Baumeister, Rubén Fernández-Busnadiego, Antonio Martínez-Sánchez

**Affiliations:** Department of Molecular Structural Biology, Max Planck Institute of Biochemistry, 82152 Martinsried, Germany; Graduate School of Quantitative Biosciences Munich, 81337 Munich, Germany

## Abstract

Curvature is an important morphological descriptor of cellular membranes. Cryo-electron tomography (cryo-ET) is particularly well-suited to visualize and analyze membrane morphology in a close-to-native state and high resolution. However, current curvature estimation methods cannot be applied directly to membrane segmentations in cryo-ET. Additionally, a reliable estimation requires to cope with quantization noise. Here, we developed and implemented a method for membrane curvature estimation from tomogram segmentations.

From a membrane segmentation, a signed surface (triangle mesh) is first extracted. The triangle mesh is then represented by a graph (vertices and edges), which facilitates finding neighboring triangles and the calculation of geodesic distances necessary for local curvature estimation. Here, we present several approaches for accurate curvature estimation based on tensor voting. Beside curvatures, these methods also provide robust estimations of surface normals and principal directions.

We tested the different methods on benchmark surfaces with known curvature, demonstrating the validity of these methods and their robustness to quantization noise. We also applied two of these approaches to biological cryo-ET data. The results allowed us to determine the best approach to estimate membrane curvature in cellular cryo-ET data.

## 1 Introduction

Membranes define the limits of the cells and encompass compartments within eukaryotic cells, helping to maintain specific micro-environments with different shapes and functions. Membrane curvature is important for many cellular processes, including organelle shaping, endo-and exocytosis, vesicle formation, scission and fusion, protein sorting and enzyme activation [McMahon and Boucrot, 2015, Bassereau et al., 2018]. There is a plethora of cellular mechanisms for generation, sensing and maintenance of local membrane curvature, e.g. clustering of conical lipids or transmembrane proteins, insertion or interaction with specific protein domains, as well as larger scale scaffolding of proteins or cytoskeletal filaments [Kozlov et al., 2014, McMahon and Boucrot, 2015]. Interestingly, membrane curvature can also have pathological implications: polyQ-expanded huntingtin exon I fibrils induce high curvature in the endoplasmic reticulum (ER) membrane, perhaps leading to ER membrane disruption [Bäuerlein et al., 2017]. Cryo-ET enables an accurate three-dimensional (3D) visualization and analysis of the subcellular architecture at high resolution [Lucic et al., 2005, Beck and Baumeister, 2016, Wagner et al., 2017] and is particularly well-suited to study membrane morphology. While other TEM techniques often cause membrane deformations, cryo-ET allows high-resolution imaging of fully hydrated biological material, by which membranes can be observed in a close-to-native state [Collado and Fernández-Busnadiego, 2017]. In the electron microscope, the sample is tilted around an axis while 2D images of a cellular region of interest are acquired for each tilt. The tilt series are then computationally aligned and reconstructed into a tomogram, which is a 3D gray-value image of the cellular interior. Because it is technically impossible to tilt the sample by 360°, there is missing information in the tomogram. This artifact, called *missing wedge* [Lucic et al., 2005], causes the membranes to look smeared out or opened in the Z-axis. Moreover in cryo-ET, the cells are illuminated by only a low dose of electrons, resulting in images of low signal-to-noise ratio. Segmentation, i.e. tracking or marking, of structural components present in tomograms is necessary for their interpretation. Available software packages can assist membrane segmentation [Martinez-Sanchez et al., 2014], but in most of the cases human supervision is still necessary due to the complexity of the cellular context and the low signal-to-noise ratio.

Currently, membrane segmentation interpretation is highly limited because of the lack of computational methods to measure quantitative descriptors. A membrane can be modeled as a 2D-manifold surface [Martinez-Sanchez et al., 2011], so curvature descriptors characterize their local geometry. The goal of this study is to quantitatively determine local curvature descriptors, i.e. principal curvatures and principal directions, of cellular membranes from tomogram segmentations. A membrane surface is best described by an oriented triangle mesh. However, triangle mesh generation from a set of voxels [Hoppe et al., 1992] is not trivial because of the presence of holes in membrane segmentations. There are many algorithms described for local curvature computation, e.g. [Taubin, 1995, Meyer et al., 2003, Theisel et al., 2004, Rusinkiewicz, 2004, Razdan and Bae, 2005, Szilvási-Nagy, 2008], and those based on tensor voting [Tang and Medioni, 1999, Page et al., 2002, Tong and Tang, 2005] have proved particularly robust against noise. Besides the errors generated during membrane segmentation, quantization noise is the limiting factor for describing local membrane geometry. Here we use the term quantization noise to refer the discretizing effect attributed to the distance between two adjacent voxels. It defines the maximum precision for sampling a membrane segmentation. Consequently, the surface curvature estimation algorithm must be adapted to leverage up the maximum precision provided by cryo-ET.

Here, we developed, implemented and compared several methods for robust membrane curvature estimation from tomogram segmentations. In brief, the workflow has the following steps: From a segmentation, a single-layered, signed triangle mesh surface is firstly extracted. The surface is then converted to a graph, representing triangle center coordinates by vertices and connecting those centers originating from adjacent triangles by edges. Finally, local curvature descriptors are computed for every triangle center. Our proposal combines two established algorithms [Page et al., 2002, Tong and Tang, 2005] based on tensor voting [Medioni et al., 2000], to increase the robustness to membrane geometries present in cryo-ET and to minimize the impact of quantization noise.

## 2 Method

### 2.1 Cryo-ET Data Collection

The *in situ* cryo-ET data used in this study was collected from vitrified yeast (*Saccharomyces cerevisiae*) cells, milled down to 150-250 nm thick lamellas using cryo-focused ion beam (FIB) [Rigort et al., 2012] and imaged using a Titan Krios cryo-electron microscope (FEI), equipped with a K2 Summit direct electron detector (Gatan), operated in dose fractionation mode. Tilt series were recorded using SerialEM software [Mastronarde, 2005] at a magnification of 42,000 X (pixel size of 3.42 Å), typically from −46° to +64° with increments of 2°. The K2 frames were aligned using K2Align software [Li et al., 2013]. Tilt series were aligned using patch-tracking and weighted back projection provided by the IMOD software package [Kremer et al., 1996]. Tomograms were binned 4 X to improve contrast prior to segmentation. The voxel size of the final segmentations was 1.368 nm. Membrane segmentations were generated automatically from tomograms using TomoSegMemTV [Martinez-Sanchez et al., 2014] and further refined manually by an expert using Amira (FEI Visualization Sciences Group). The lumen of membrane compartments was then filled manually.

### 2.2 Surface Generation

#### Membrane Segmentations

The segmented membrane of interest (Fig. 1B) is used as the input for a surface reconstruction algorithm that reconstructs closed surfaces without boundaries from unorganized set of points [Hoppe et al., 1992], here the membrane voxels (Fig. 2A). However, most segmented membranes in cryo-ET are open, e.g. due to missing wedge artifacts. Attempting to close the surface, the algorithm generates surface regions outside the segmented mask. Therefore, we discard these regions by applying a mask using the membrane segmentation.

**Figure 1:**
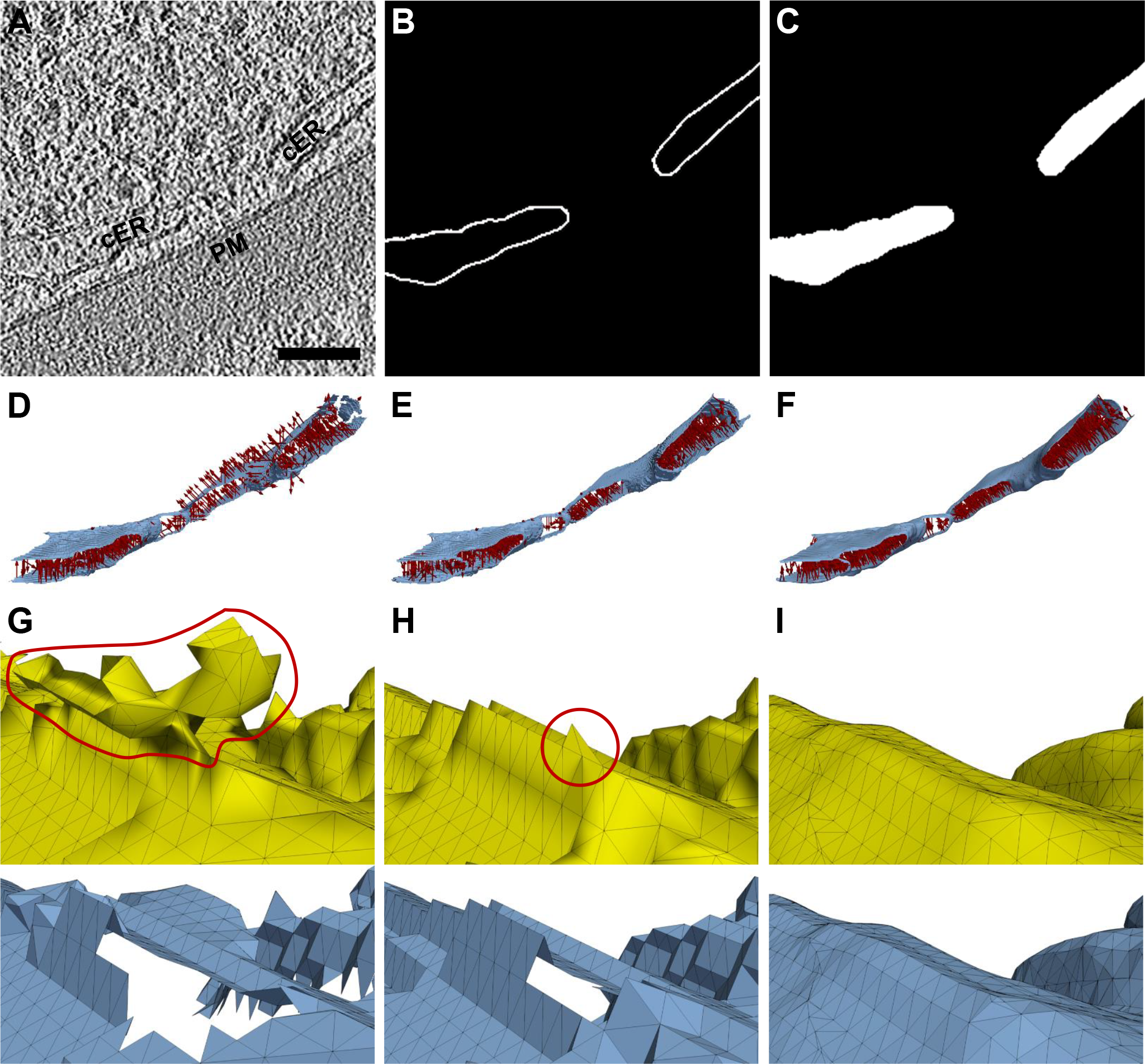
Comparison of surface generation methods. (**A**) A tomographic slice showing the cortical ER (cER) and plasma membrane (PM) of a yeast cell (scale bar: 100 nm). Panels (B-C) show the same slice as in (A) with (**B**) the membrane segmentation and (**C**) the compartment segmentation of the cER membrane. Panels (D-F) Surface normals (every 100th) are shown as dark red arrows. (**D**) A surface generated from the cER membrane segmentation shown in (B); some normals erroneously point outside the cER lumen. Panels (E-F) show surfaces generated using the compartment segmentation shown in (C), (**E**) without and (**F**) with Gaussian smoothing; all normals point inside the cER lumen. Panels (G-I) show a different magnified view of the surfaces shown in panels (D-F) in the same order. The top surfaces in yellow are simply masked by the membrane segmentation. The bottom surfaces in blue are additionally cleaned from artifact borders using the graph. (**G**) A red line marks an artifact in the masked surface leading to a hole in the clean surface. (**H**) Sticking out triangles (inside the red circle) in the masked surface leading to a hole in the clean surface. (**I**) The smoothed surface is free from artifacts.

**Figure 2:**
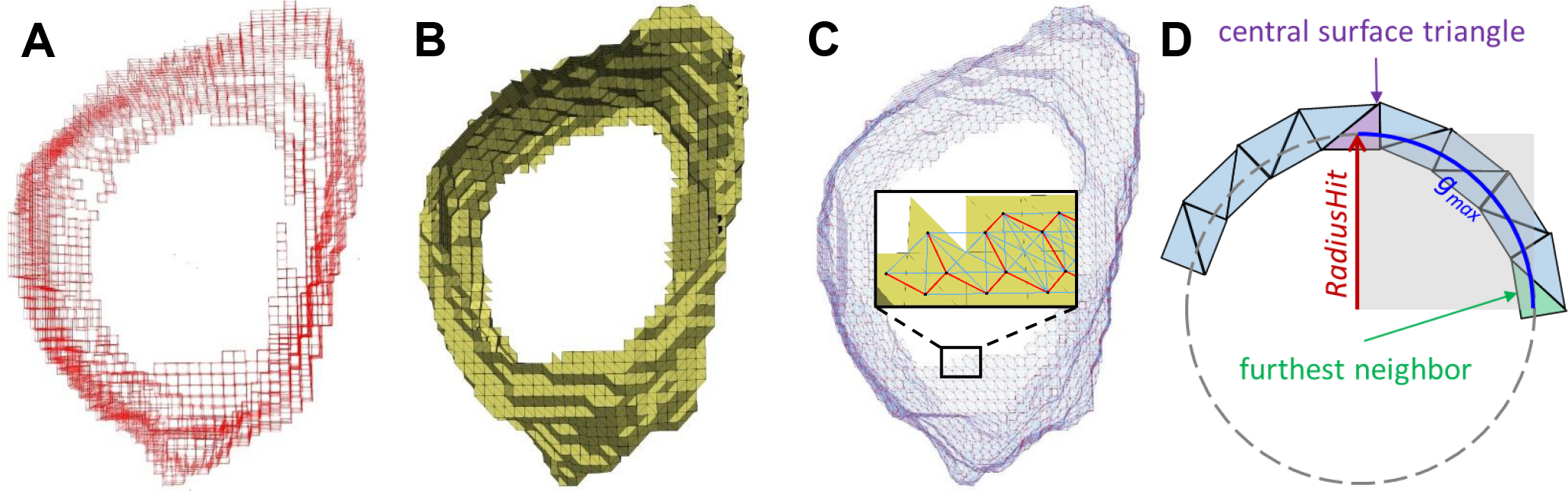
Surface Graph generation and parameters defining the geodesic neighborhood. (**A**) Voxels of a segmentation of a vesicle from a cryo-electron tomogram ([Bäuerlein et al., 2017]). (**B**) A surface (triangle mesh) generated from the membrane segmentation shown in (A). (**C**) Surface Graph generated from the surface shown in (B); the inset shows a magnified region of the graph mapped on top of the triangle mesh (triangles: yellow, graph vertices: black dots, strong edges: red, weak edges: light blue). (**D**) Schematic illustrating *RadiusHit* and *g*_*max*_ parameters. *g*_*max*_ is ¼ of the circle perimeter with radius equal to *RadiusHit*. *g*_*max*_ defines the maximal geodesic distance from a surface triangle center to the centers of its neighboring triangles, approximated by the shortest path along the edges of the Surface Graph.

We use the convention that normals should point inwards in a convex surface. However since membrane segmentations have boundaries, the algorithm [Hoppe et al., 1992] sometimes mistakenly initiates normals on both sides (Fig. 1D). As a result, ridge-like patches appear along the surface, leading to holes in the clean surface (Fig. 1G). In some cases the surface reconstruction is improved by closing small holes in the segmentation using morphological operators.

#### Compartment Segmentations

This surface generation method requires additional segmentation of the inner volume of a compartment enclosed by a membrane (Fig. 1C). This unequivocally defines the orientation of the membrane by closing its holes. This process is usually more time consuming because of the lack of computational methods to assist this task. However, surface orientation is recovered perfectly (Fig. 1E). After joining the membrane and its inner volume masks, we generate an isosurface around the resulting volume using the Marching Cubes algorithm [Lorensen and Cline, 1987]. Finally, we apply a mask using the original membrane segmentation to keep only the surface region going through the membrane.

In some cases, especially where the membrane segmentation was manually refined, Marching Cubes produces triangles standing out perpendicularly to the surface (Fig. 1H, top). To correct those artifacts and exploit the subvoxel precision offered by Marching Cubes, the compartment segmentation mask was slightly smoothed using a Gaussian kernel with *σ* = 1 voxel before extracting the surface (Fig. 1F, I).

Thus, we have two procedures for surface generation from segmentations: The first procedure can work directly on membrane segmentations but the output surfaces tend to contain artifacts, whereas the second one ensures smoother and well oriented surfaces but requires more human intervention. As mentioned, in both procedures we apply a mask on the generated surface using the membrane segmentation to keep only the surface region that goes through the membrane. A distance of 3 pixels to the mask is used to bridge upon small holes in the segmentation. However, this also results in additional surface borders of three pixels in width. Those borders have to be removed before curvature calculation, as described next.

### 2.3 Surface Graph Generation

Curvature is a local property that has to be estimated using a local neighborhood of triangles. If the neighborhood is too small, we would measure only noise created by the steps between voxels. If the neighborhood is too large, we would underestimate the curvature. Some existing approaches do not consider a sufficient geodesic neighborhood for curvature estimation (e.g. in VTK library [Schroeder et al., 2006]).

To accurately estimate geodesic distances, we used the graph-tool python library [P. Peixoto, 2014] to map the triangle mesh (Fig. 2B) into a spatially embedded graph, here referred as Surface Graph. First, graph vertices are associated to triangle centroid coordinates. Second, pairs of graph vertices representing triangles sharing two triangle vertices are connected by *strong* edges, while those sharing only one triangle vertex are connected by *weak* edges (Fig. 2C). Geodesic distances are computed from the shortest paths found between triangle centers along the graph edges [Dijkstra, 1959]. Here, both strong and weak edges are used for a better approximation. Using the graph, we can detect triangles at surface borders because they have less than three strong edges, and then find triangles up to a certain geodesic distance from the border triangles and filter them out from the surface.

Now that we have a surface of a membrane and a tool to find the geodesic neighborhood and discard surface borders artifacts, we can proceed with curvature estimation.

### 2.4 Curvature Estimation

Our algorithm combines two previously published algorithms that are based on tensor voting and curvature tensor theory [Page et al., 2002, Tong and Tang, 2005], to increase the precision of curvature estimation for our surfaces. In order to estimate principal curvatures, principal directions have to be estimated. For the estimation of principal directions, surface normals are required. The true surface normals are estimated by averaging normals of triangles within a geodesic neighborhood.

#### 2.4.1 Parameters Defining the Geodesic Neighborhood

Similarly to [Tong and Tang, 2005], here we define a *RadiusHit* parameter to approximate the highest curvature value we can estimate reliably, which is 1/*RadiusHit*. For each surface triangle center, we define its local neighborhood as

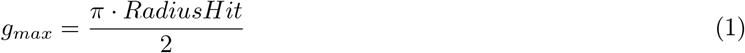

where *g*_*max*_ defines the maximal geodesic distance. In Eq. 1, *g*_*max*_ is approximated by ¼ of a circle perimeter with radius equal to *RadiusHit* (Fig. 2D).

#### 2.4.2 Robust Estimation of Surface Normals

Normals computed directly from triangle mesh are corrupted by quantization noise. To avoid this, we have adapted the first step of normal vector voting algorithm as proposed in [Page et al., 2002], but estimating the normals for each triangle center instead of defining new normals at each triangle vertex.

##### Collection of Normal Votes

For each triangle centroid (or graph vertex) v, the normal votes of all triangles within its geodesic neighborhood are collected and the weighted covariance matrix sum *V*_v_ is calculated. More precisely, a normal vote 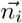 of a neighboring triangle (whose center c_*i*_ is lying within *g*_*max*_ of vertex v) is calculated using the normal 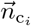 assigned to this triangle:

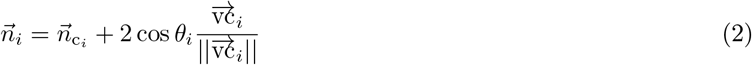

where 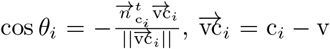. and 0 ≤ *θ_i_* ≤ *π*. This formula fits a smooth curve from c_*i*_ to v, allowing the normal vote 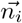 to follow this curve. According to the perceptual continuity constrain [Medioni et al., 2000], the most appropriate curve is the shortest circular arc.

Then, each vote is represented by a covariance matrix 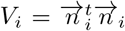 and votes from the geodesic neighborhood are collected as a weighted matrix sum *V*_v_:

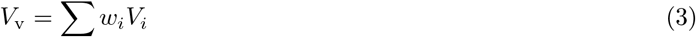

where *w*_*i*_ is a weighting term calculated as follows:

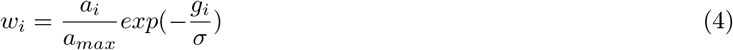

Vote weight of a neighboring triangle increases linearly with its surface area *a*_*i*_, but decreases exponentially with its geodesic distance *g*_*i*_ to v. *a*_*max*_ is the area of the largest triangle in the whole surface and *σ* is an exponential decay parameter, which is set to fulfill 3*σ* = *g*_*max*_, so that votes beyond the geodesic neighborhood have almost no influence and can be ignored.

##### Estimation of Normal Vectors

The votes collected into the matrix *V*_v_ are used for estimating the correct normal vector for the triangle represented by vertex v. This is done by eigen-decomposition of *V*_v_, which generates three real eigenvalues *e*_1_ ≥ *e*_2_ ≥ *e*_3_ with corresponding eigenvectors 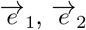 and 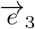. Then, the estimated normal direction is equal in its absolute value to that of the first eigenvector.

During construction of *V*_v_, the sign of normal votes is lost when *V*_*i*_ is computed. The correct orientation can be recovered from the original normal 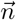, as the original surface was already oriented. Therefore, the estimated normal is correctly oriented by:

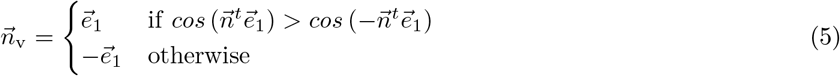

#### 2.4.3 Estimation of Principal Directions and Curvatures

For each graph vertex v, we use the estimated normals 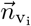 of its geodesic neighbors v_*i*_ in order to cast curvature votes. The curvature votes are summed into a curvature tensor. The resulting curvature tensor is decomposed to find the principal directions and curvatures at vertex v. Here we describe the basic curvature estimation algorithm as an adaptation of [Page et al., 2002].

#### Collection of Curvature Votes

Each neighboring vertex v_*i*_ casts a vote to the central vertex v, where the votes are collected into a 3×3 symmetric matrix *B*_v_ [Taubin, 1995]:

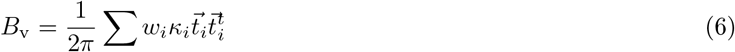

For each v_*i*_, three variables are computed:

1. Weight *w*_*i*_ depending on the geodesic distance between v_*i*_ and v, as defined in Eq. 4 but without normalizing by relative triangle area:

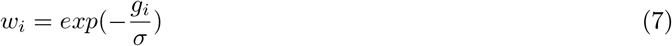

Also, all weights around the vertex v are constrained by ∑*w*_*i*_ = 2*π*.
2. Tangent 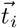 from v in the direction of the arc connecting v and v_*i*_ (using the estimated normal 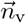 at v):

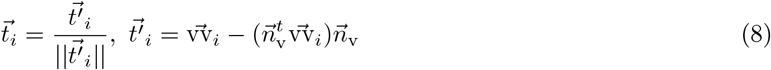
3. Normal curvature *κ_i_* [Tong and Tang, 2005]:

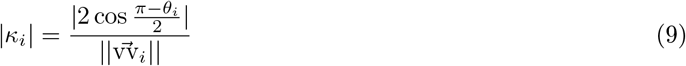

where *θ*_*i*_ is the turning angle between 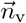 and the projection 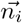 of 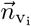 onto the arc plane (formed by v, 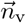 and v_*i*_). The following calculations lead to *θ*_*i*_:

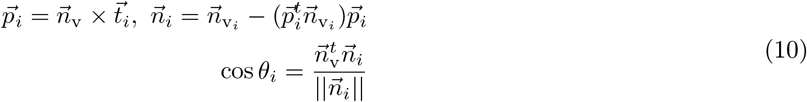

For surface generation, we use the convention that normals point inwards in a convex surface. Then, the curvature is positive if the surface patch is curved towards the normal and negative otherwise. Therefore, the sign of *κ*_*i*_ is set by:

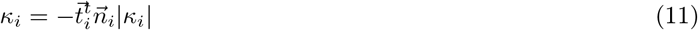

##### Curvature Estimation

For a vertex v and its calculated matrix *B*_v_, we calculate the principal directions, 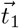 and 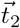, and curvatures, *κ*_1_ and *κ*_2_, at this vertex. This is done using eigen-decomposition of *B*_v_, resulting in three eigenvalues *b*_1_ ⩾ *b*_2_ ⩾ *b*_3_ and their corresponding eigenvectors 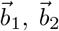 and 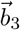. The eigenvectors 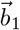 and 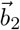 are the principal directions. The principal curvatures are found with linear transformations of those eigenvalues [Taubin, 1995]:

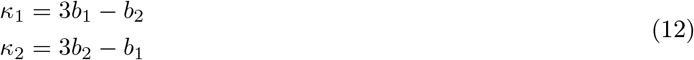

The smallest eigenvalue *b*_3_ has to be close to zero and the corresponding eigenvector 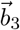 has to be similar to the normal 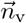 [Page et al., 2002].

#### 2.4.4 Method Variants

Besides the method described above, we have also implemented some modifications to determine the best solution for cryo-ET. Results with the comparison of the different methods with both synthetic and experimental data are shown in Sec. 3. Here is an overview of the compared methods.

##### Regular Vector Voting (RVV)

Refers to the main algorithm described above.

##### Normal Vector Voting

In [Page et al., 2002], curvature is computed as the turning angle *θ*_*i*_ divided by arc length between the vertices v and v_*i*_, which is the geodesic distance between them, *g*_*i*_:

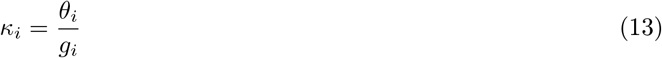

However, this definition of *κ_i_* with the sign according to our normals convention (Eq. 11) lead to erroneous eigenvalue analysis of *B*_v_. The eigenvalue analysis was only successful for *κ*_1_ *>* 0, leading to wrong curvature sign estimation for non-convex surfaces. The wrong and corrected curvature sign estimation are shown in Sec. 3.1.3.

##### Augmented Vector Voting (AVV)

Here, the weights of curvature votes, prioritizing neighbors with a closer geodesic distance to the central triangle vertex v, are normalized by relative triangle area as for normal votes using Eq. 4 instead of Eq. 7.

##### Surface Sampling Vector Voting (SSVV)

We implemented the algorithm "GenCurvVote" from [Tong and Tang, 2005] to estimate the principal directions and curvature (Sec. 2.4.3). The main difference between SSVV and RVV is that instead of using all points within the geodesic neighborhood of a given surface point v, in SSVV only eight points on the surface are sampled using *RadiusHit*. For this, an arbitrary tangent vector at v with length equal to *RadiusHit* is first generated, creating a point v_*t*_ in the plane formed by this tangent and the normal at v, 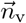. Then, a line at v_*t*_ and parallel to the normal 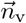 is drawn and its intersection point c with the surface is found. The tangent is rotated seven times around the normal by *π/*4 radians and same procedure is applied, leading to eight intersection points. Each vote is weighted equally, thus Eq. 6 simplifies to:

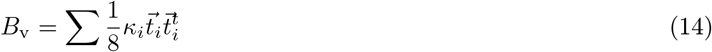

The output of our methods comprises the surface with corrected normals, estimated principal directions and curvatures as well as different combined curvature measures: mean curvature, Gauss curvature, curvedness and shape index [Koenderink and van Doorn, 1992].

##### VTK

We also calculated surface curvature using the VTK library [Schroeder et al., 2006]. VTK calculates curvature per triangle vertex using only its adjacent triangles and applying discrete differential operators [Meyer et al., 2003]. In order to be able to compare VTK to our tensor voting-based methods operating on triangles, we average the values of each curvature type at three triangle vertices to obtain one value per triangle. VTK does not estimate principal directions.

## 3 Evaluation Results

In the first part of this section, we explain how we evaluate our curvature estimation methods on several benchmark surfaces and show the quantitative results. In the second part, we present results of the best-performing methods on real biological membranes from cryo-ET data, and give recommendations regarding the choice of *RadiusHit* and borders filtering parameters.

### 3.1 Quantitative Results on Benchmark Surfaces

#### 3.1.1 Calculation of Errors

To evaluate the accuracy of our method using benchmark surfaces with known orientation and curvature, we use two following types of errors:

1. For vectors (normals or principal directions):

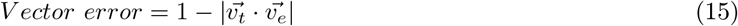

where 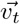 is a true vector and 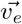 is an estimated vector for the same triangle. The minimum error is 0, when the true and estimated vectors are parallel, and the maximum error is 1, when the vectors are perpendicular. Estimated normals and principal directions should not be more than 90° misoriented since the surface is signed.
2. For scalars (principal curvatures) we use:

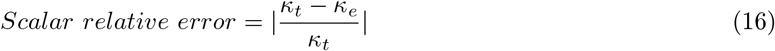

where *κ*_*t*_ is a true curvature and *κ_e_* is an estimated curvature for the same triangle. The minimum error is 0, when the estimate equals to the true value, and there is no upper bound to the error.

#### 3.1.2 Robust Estimation of Normals

Surface normals are required for a reliable estimation of the principal directions, for which the principal curvatures are estimated. In this experiment, we wanted to check that the first step of the algorithm (Sec. 2.4.2), which is the same for all our tensor voting-based methods and is thus called Vector Voting (VV) here, restores the correct orientation of the normals. For this, we use a plane surface with artificially introduced noise to simulate the quantization noise present in surfaces generated from cryo-ET segmentations. The true normals are those from the plane without noise (i.e. parallel to Z-axis, Fig. 3A). Noise was introduced to the original plane by moving each triangle vertex in the direction of its normal vector with Gaussian variance equal to 10% of the average triangle edge. As a result, the triangle normals of the noisy plane are not parallel to each other and to Z-axis (Fig. 3B, E). Using VV with *RadiusHit* of 4 pixels, the original normal orientation is almost restored (Fig. 3C, E). Using *RadiusHit* of 8 pixels, the estimation becomes even better (Fig. 3D, E), since more neighboring triangles help to average out the noise. Thus, the first step of our methods with a high enough *RadiusHit* restores the original surface direction.

**Figure 3:**
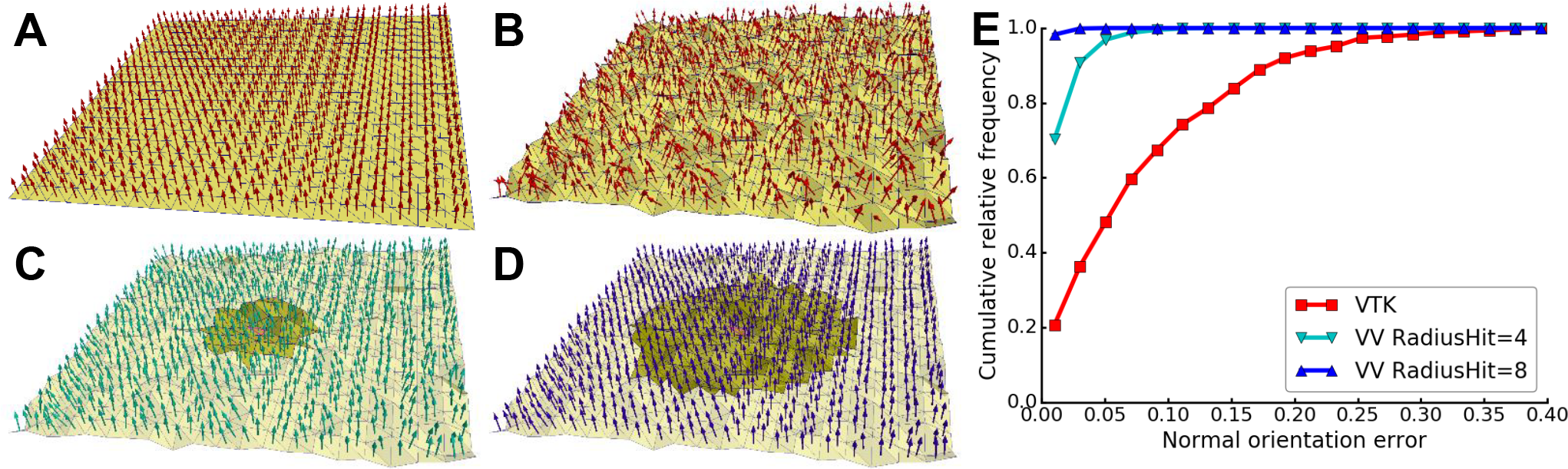
Estimation of normals on a noisy plane. (**A**) True normals (red arrows) on a smooth plane surface (yellow). (**B**) Normals on a noisy (10%) plane, calculated by VTK. Panels (C-D) show normals on the noisy plane corrected by VV with *RadiusHit* of (**C**) 4 or (**D**) 8 pixels. The neighborhoods of a central triangle are shown in a darker color. (**E**) Cumulative relative frequency histogram of normal orientation error for the noisy (10%) plane: uncorrected (VTK), corrected by the first step of our tensor voting-based methods (VV) with *RadiusHit* of 4 or 8 pixels. Curve colors in (E) correspond to the colors of the normals in (B-D).

#### 3.1.3 Estimation of the Curvature Sign

To determine the correct procedure for curvature sign determination, we used a torus as a benchmark, as this surface has regions with both positive, *κ*_1_*κ*_2_ *>* 0, and negative, *κ*_1_*κ*_2_ *<* 0, Gaussian curvature (*κ*_2_ is shown in Fig. 4A). Whereas NVV [Page et al., 2002] does not distinguish negative from positive regions (Fig. 4B), RVV and SSVV differentiate these regions correctly (Fig. 4C, D).

**Figure 4:**
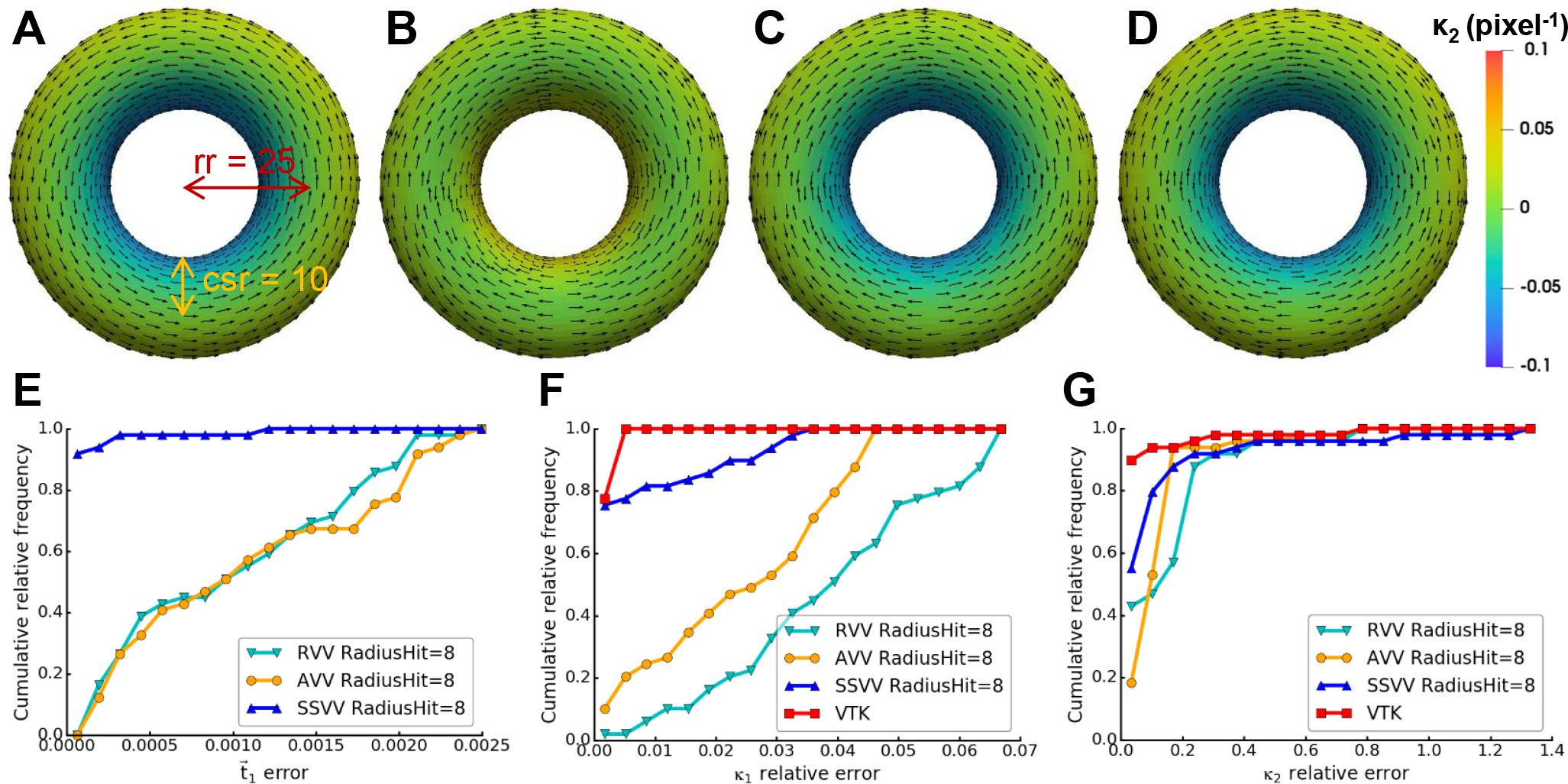
Curvature sign, principal direction and curvature estimation accuracy for a torus. Panels (A-D) show visualizations of *κ*_2_ (pixel^−1^, triangles are color-coded by curvature, see scale bar) and 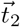 (every fourth vector is shown as an arrow from a triangle center): (**A**) true values calculated analytically for a smooth torus surface with ring radius (rr)=25 pixels and cross-section radius (csr)=10 pixels, (**B**) estimated values using NVV, (**C**) RVV or (**D**) SSVV, in all cases using *RadiusHit* = 8. Panels (E-G) show cumulative relative frequency histograms of the principal direction and curvature errors: (**E**) 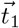 (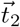 not shown as it is very similar), (**F**) *κ*_1_, (**G**) *κ*_2_ on the torus surface using different methods: RVV, AVV and SSVV (*RadiusHit*=8 pixels for all) and VTK (only for curvatures).

Thus, our RVV and SSVV methods correct the curvature sign estimation of the original method, NVV. Since RVV and SSVV both calculate the normal curvature using Eq. 9 and 11, while NVV uses Eq. 13, the latter must be the source of the erroneous curvature sign estimation. Therefore, we exclude NVV from our consideration and further comparisons.

#### 3.1.4 Setting the RadiusHit Parameter

As shown in Sec. 3.1.2, neighborhood size defined by the *RadiusHit* parameter influences the estimation of normals. Therefore, choosing an appropriate *RadiusHit* for the data is crucial for accurate curvature estimation.

To study the influence of *RadiusHit* in our different curvature estimation methods, we generated a synthetic segmentation (25³ voxels) of a sphere with radius of 10 pixels, emulating the quantization noise present in cryo-ET data (Fig. 5A). Then, we generated an isosurface of this segmentation and estimated its curvature using the different methods and *RadiusHit* values. Tong and Tang suggested that the optimal *RadiusHit* leads to the sharpest peak in the histograms around the true curvature value [Tong and Tang, 2005]. For this sphere surface, the sharpest peak close to the true curvature value, 1/10 = 0.1 pixels^−1^, is reached for *RadiusHit*=9 pixels for RVV, AVV and SSVV (Fig. 5B-D). This value is close to the sphere radius, indicating that the most robust estimation can be achieved using a *RadiusHit* approximately similar to the feature radius. Moreover, the peaks get sharper with higher *RadiusHit* values, i.e. there are less wrongly assigned values. This is consistent with the fact that the larger the neighborhood the higher the robustness against noise. On the other hand, features with a radius less than *RadiusHit* are averaged (RVV and AVV) or are neglected (SSVV), so 1*/RadiusHit* can be set as a limit for the maximal curvature that can be reliably computed.

**Figure 5:**
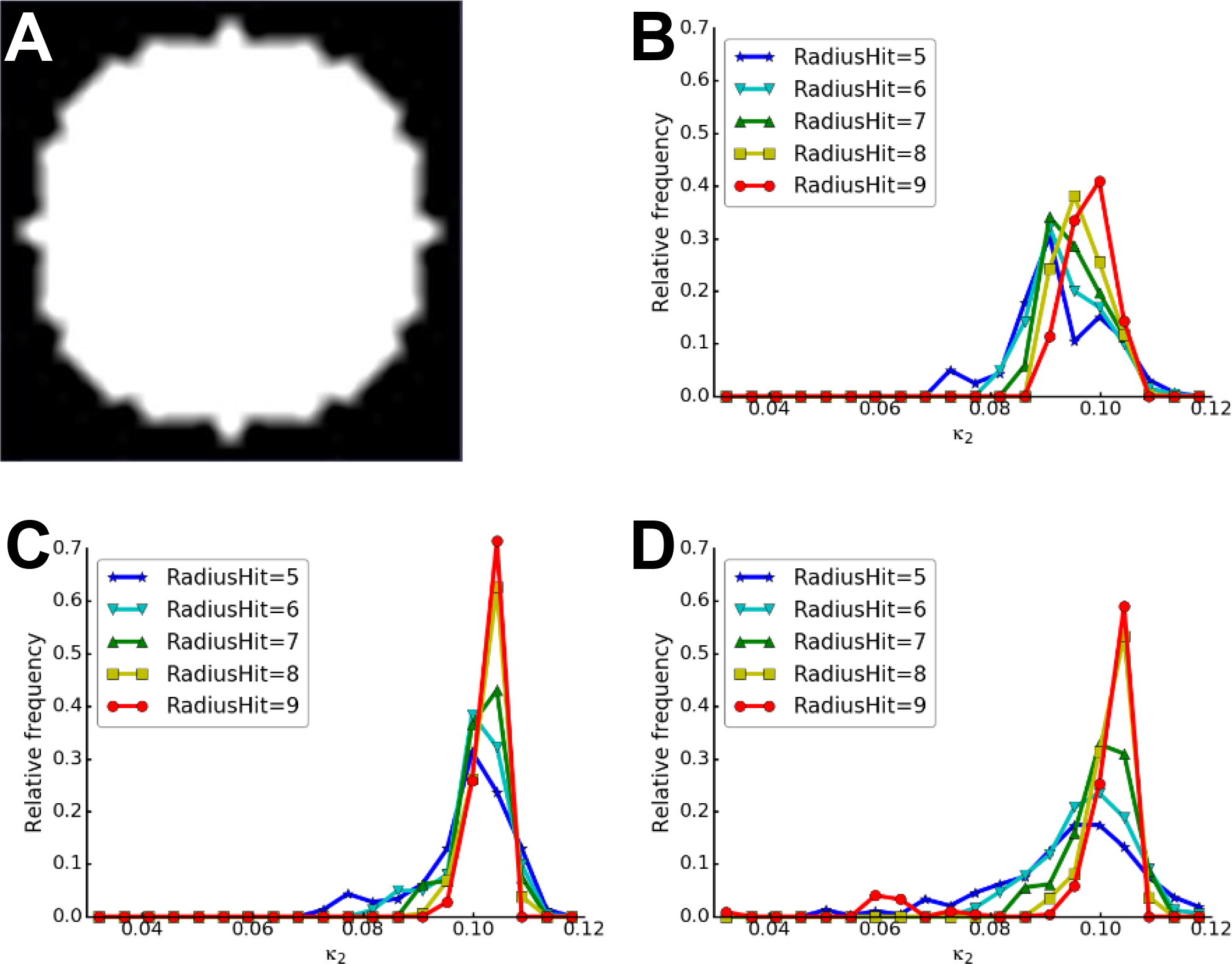
RadiusHit parameter choice. (**A**) A central slice of a synthetic segmentation of a sphere with radius=10 pixels. Panels (B-D) show histograms of the estimated *κ*_2_ on the isosurface extracted from the segmentation shown in (A), using different *RadiusHit* (5-9) and methods: (**B**) RVV, (**C**) AVV and (**D**) SSVV.

#### 3.1.5 Estimation Accuracy on Smooth Surfaces

To evaluate the performance of the different curvature estimation methods, we first calculated the principal directions and curvatures errors on smooth surfaces.

For the smooth torus surface shown in Fig. 4A-D, SSVV estimates both principal directions (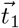 error histogram is shown in Fig. 4E, very similar results for 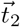 are not shown) and curvatures (Fig. 4F-G) more accurately than RVV, but the error difference is very small. AVV slightly outperforms RVV in the estimation of principal curvatures for this surface. However, VTK estimates principal curvatures a bit better than the tensor voting-based methods for this very smooth surface with uniform triangles. Note that the curvature errors for *κ*_1_ (Fig. 4F) are lower than for *κ*_2_ (Fig. 4G) for all methods. A possible explanation is that *κ*_1_ is constant for torus (reciprocal to the cross-section radius) and thus easier to estimate, while *κ*_2_ is changing depending on the position on a cross-section circle.

We also compared the methods using a smooth sphere surface with a non-uniform triangle tessellation, generated within a volume using a 3D Gaussian function (*σ*=3.3) and applying an isosurface. Since both principal curvatures should be the same for a sphere surface, they were considered together. Also, no true principal directions exist for a sphere surface. For a sphere with radius=10 pixels, a *RadiusHit* value of 9 pixels was used for all methods. VTK has very high errors (maximal error~8, Fig. 6A). The maximal error is only 0.18 for RVV (Fig. 6B, D), while AVV achieves a substantial improvement (maximal error~0.02) over RVV (Fig. 6C, D). Presumably because of the non-uniform tessellation (Fig. 6A-C), weighting by triangle area matters more for this surface, so the performance improvement of AVV over RVV is more evident here than for the uniformly tessellated torus surface. SSVV performs slightly better than AVV (Fig. 6D).

**Figure 6:**
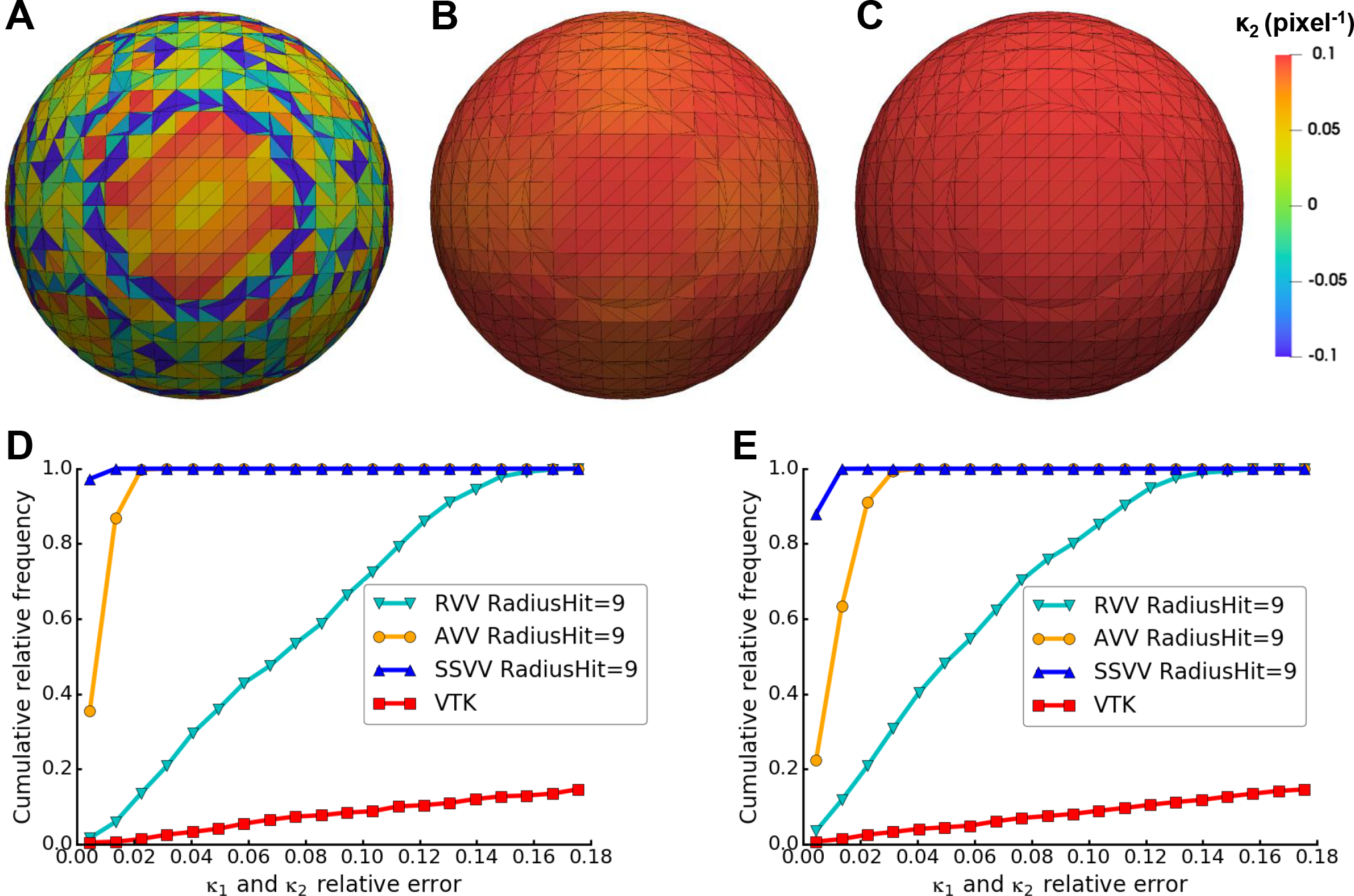
Accuracy of curvature estimation on a smooth spherical surface. Panels (A-C) show visualizations of *κ*_2_ (pixel^−1^) estimated by (**A**) VTK, (**B**) RVV or (**C**) AVV on a smooth sphere with radius=10 pixels. Panels (D-E) show cumulative relative frequency histograms of the principal curvature (*κ*_1_ and *κ*_2_ taken together) errors on a smooth sphere with radius=10 (**D**) or 20 pixels (**E**) using different methods: RVV, AVV and SSVV (*RadiusHit*=8 pixels) and VTK.

To test how stable the methods are for different curvature scales, we increased the radius of the smooth sphere from 10 to 20 pixels, while leaving *RadiusHit* the same (9 pixels). As a result, the errors increase drastically for VTK (maximal error ~18), while the tensor voting-based methods perform almost the same as for the sphere with radius 10 pixels (Fig. 6E). Similar results were obtained for a sphere with radius=30 voxels (not shown).

Altogether, the evaluation results on smooth benchmark surfaces show that the tensor voting-based methods are quite stable to feature sizes variations (beyond *RadiusHit*) and irregular triangles within one surface, especially SSVV, whereas VTK only performs well for a very smooth surface with a regular triangulation. AVV can deal with non-uniformly tessellated surfaces better than RVV. This is because curvature votes are also weighted by relative triangle area in AVV. In the original algorithm [Page et al., 2002], weighting curvature votes by triangle area would not make sense because normals and curvature are estimated at triangle vertices. Since we decided to estimate normals and curvature at triangle centers instead, curvature votes are cast by complete triangles and weighting them by triangle area proved to be advantageous. Therefore, we exclude RVV from further comparison.

#### 3.1.6 Robustness to Surface Noise

Until now we evaluated the performance of the methods on smooth surfaces. However, surfaces generated from membrane segmentations are not smooth, as the surface triangles tend to follow the voxel boundaries resulting in a step-like surface. Such quantization noise depends on the voxel size of the segmented tomogram.

To test how the methods perform in presence of quantization noise, we generated a step-like surface of a sphere with a radius of 10 pixels, as described in Sec. 3.1.4. As expected, VTK just measures the curvature differences between directly neighboring triangles, resulting in high errors (Fig. 7A). Therefore, VTK was excluded from the subsequent error calculations. To compare AVV and SSVV, we first used the sphere with radius=10 pixels and the optimal *RadiusHit* value of 9 pixels (Fig. 5C-D). As expected, the principal curvature errors are higher for both methods (10-20 fold, Fig. 7D) compared to the smooth sphere (Fig. 6D). However, in this case AVV outperforms SSVV (Fig. 7B-D). On the smooth spherical surface, SSVV performed slightly better than AVV (Fig. 6D).

**Figure 7:**
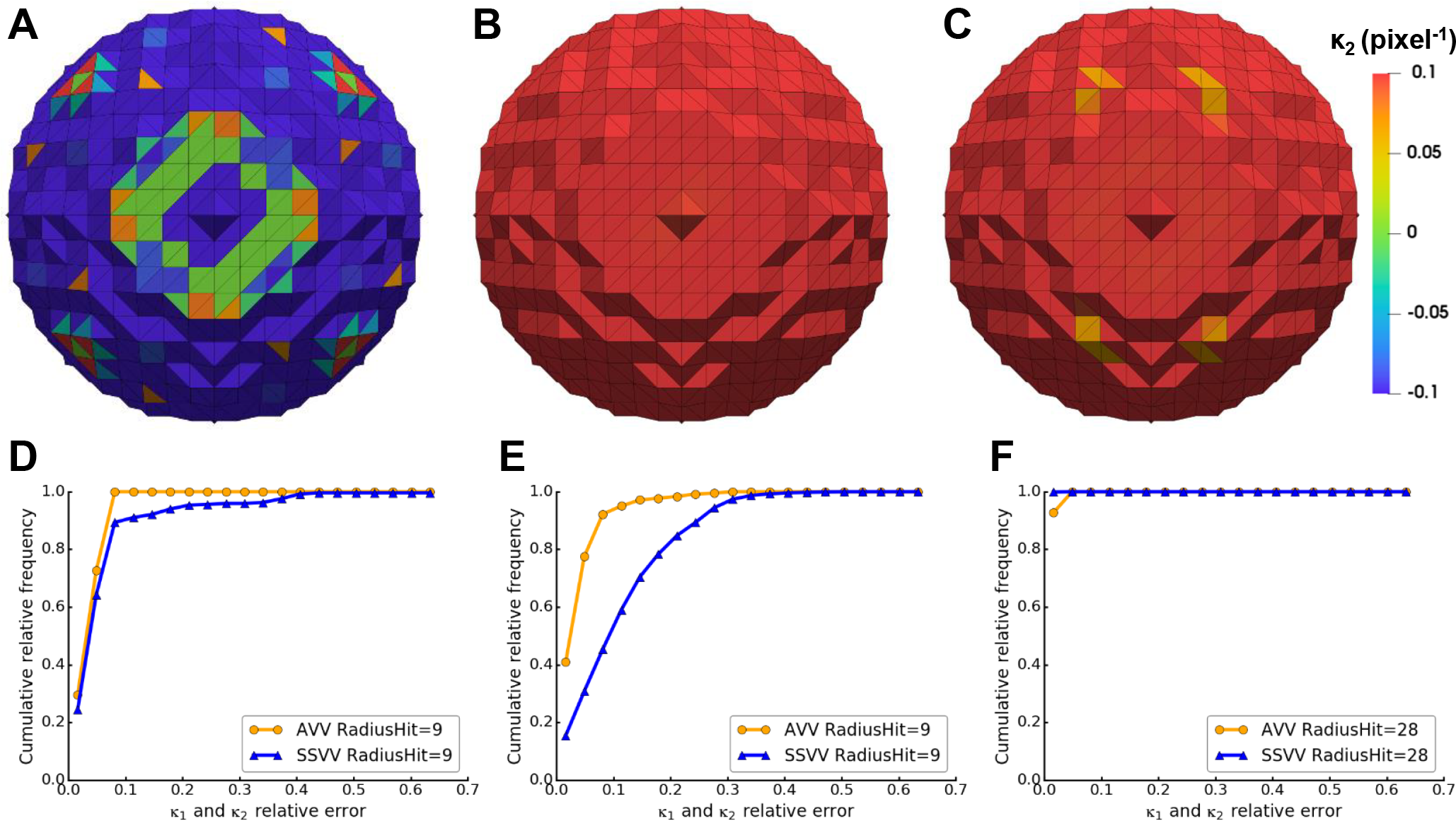
Accuracy of curvature estimation on a spherical surface with quantization noise. Panels (A-C) show visualizations of *κ*_2_ (pixel^−1^) estimated by (**A**) VTK, (**B**) AVV or (**C**) SSVV on a sphere with quantization noise and radius=10 pixels. Panels (**D**-**F**) show cumulative relative frequency histograms of the principal curvature (*κ*_1_ and *κ*_2_ taken together) errors on a spherical surface with quantization noise and radius=10 (D) or 30 voxels (E-F) using AVV and SSVV with different *RadiusHit* values.

Again, we wanted to compare how the estimation accuracy of the methods decreases with increasing the size of the feature, for a constant *RadiusHit* value. When using a sphere with radius=20 pixels, the estimation accuracy of SSVV decreases (not shown), and even more with radius=30 pixels, while the performance of AVV decreases only slightly (Fig. 7E). The performance of SSVV only improves by using a *RadiusHit* almost as high as the sphere radius (*RadiusHit*=18 or 28 for radius=20 or 30 pixels, respectively), which should be close to optimum (Fig. 7F). Note that AVV with the same *RadiusHit* also performs better and similarly to SSVV. Moreover, performance of both methods increases with the sphere size when the optimal *RadiusHit* is used (compare Fig. 7D and F).

The evaluation results on a noisy sphere surface show that when quantization noise is present, SSVV requires a higher *RadiusHit* similar to the curvature radius, while AVV is quite stable with a lower *RadiusHit* range. Using a very high *RadiusHit* is generally not advisable, as it would lead to the underestimation of the curvatures of smaller surface features. Thus, AVV is the more robust algorithm in this test.

#### 3.1.7 Higher Errors at Surface Borders

As explained with respect to surface generation (Sec. 2.2), membranes in cryo-ET segmentations have holes and open ends. We aimed for a curvature estimation algorithm that is robust to such artifacts.

Tensor voting-based methods use a supporting neighborhood in order to improve the estimation, so holes much smaller than the neighborhood region are not critical. However, a vertex close to surface border can have considerably less neighbors. Therefore, we hypothesized that the estimation at vertices close to such borders would be worse. To prove this hypothesis, we generated a smooth cylinder surface opened at both sides with radius=10 and height=25 pixels and evaluated the performance of our methods.

As expected, both tensor voting-based methods make a worse estimation near borders: AVV overestimates *κ*_1_ gradually when moving towards the border and *κ*_2_ at the very border (Fig. 8A), while SSVV underestimates *κ*_1_ consistently and makes a gradient of wrong estimations for *κ*_2_ in the same region (Fig. 8B). Since VTK does not use a bigger neighborhood, it shows high errors all over the cylinder, where triangle pattern changes (Fig. 8C). SSVV makes slightly higher errors for 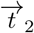 than AVV (Fig. 8D), but performs slightly better for *κ*_1_, while VTK makes higher *κ*_1_ errors than the other two methods (Fig. 8E). When excluding values within distance of 5 voxels of the graph border, the errors are virtually eliminated for AVV and SSVV, but do not change for VTK (Fig. 8F-G).

**Figure 8:**
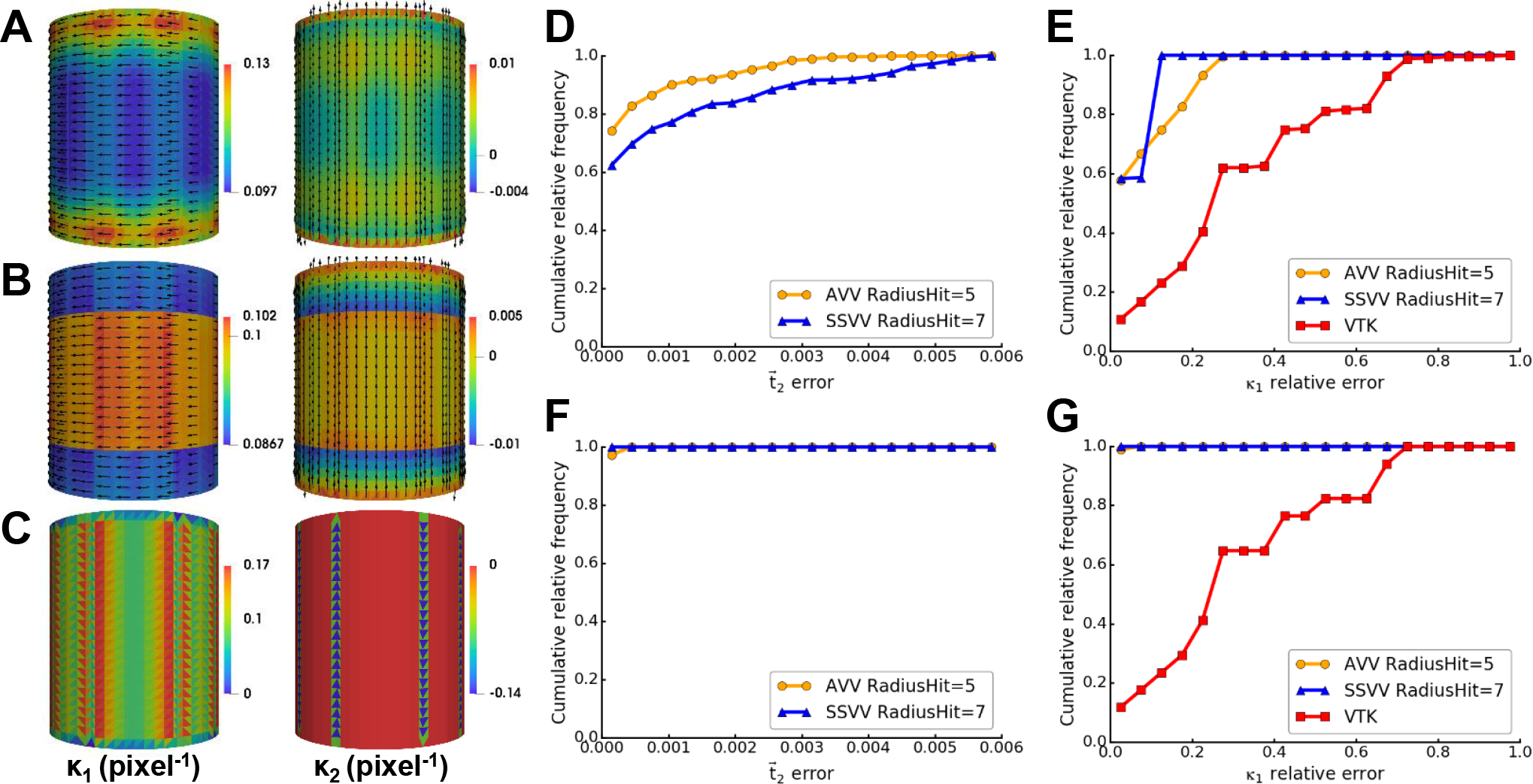
Estimation accuracy near borders of a cylinder surface. Panels (A-C) show principal curvatures on a smooth cylinder surface with radius=10 and height=25 pixels estimated by different methods: (**A**) AVV, (**B**) SSVV, (**C**) VTK; the estimated *κ*_1_ and *κ*_2_ are shown in their original range; true *κ*_1_=0.1, true *κ*_2_=0 pixels^−1^. Panels (**D**-**G**) show cumulative relative frequency histograms of the 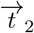 errors (D, F) and *κ*_1_ relative errors (E, G) on the cylinder surface using different methods: AVV, SSVV (using optimal *RadiusHit*=5 or 7 pixels, respectively) and VTK (only for *κ*_1_). In (F-G) values within 5 pixels of the graph border are excluded.

Altogether, the benchmark results demonstrate the validity of our tensor voting-based methods and their robustness to quantization noise of triangle meshes originating from tomogram segmentations, especially of AVV. Additionally, curvature estimations at surface borders can be erroneous, so they should be excluded from analysis.

### 3.2 Application on Biological Membranes

Lastly, we tested our methods on real biological membranes from cryo-ET data. Here, we compare performance of AVV and SSVV on surfaces generated using membrane and compartment segmentations of yeast cortical ER (cER). We also study the behavior of the methods on a complex feature for different *RadiusHit* values. Finally, we suggest a simple method to avoid edge artifacts.

#### 3.2.1 Methods and Parameter Choice

We used a cER membrane segmentation that contains several high curvature regions in the form of rounded cones or peaks to evaluate the performance of AVV and SSVV, the methods that proved most robust to quantization noise (Sec. 3.1).

First, we studied the relationship between the *RadiusHit* parameter and the feature size. For this, we isolated a single cER peak with a base radius of approximately 10 nm and estimated its curvature using several *RadiusHit* values (histograms in Fig. 9). For *RadiusHit* below 10 nm, the *κ*_1_ distributions are broader. For *RadiusHit*=10 nm, *κ*_1_ distributions are enriched in the vicinity of 0.1 nm^−1^, which means that the feature starts to be observed as a whole and its smaller components fade. By increasing *RadiusHit* parameter above the radius of the feature, the curvature distributions shift towards lower values and become sharper (especially for SSVV), underestimating the curvature of the feature, i.e. the feature is averaged out.

**Figure 9:**
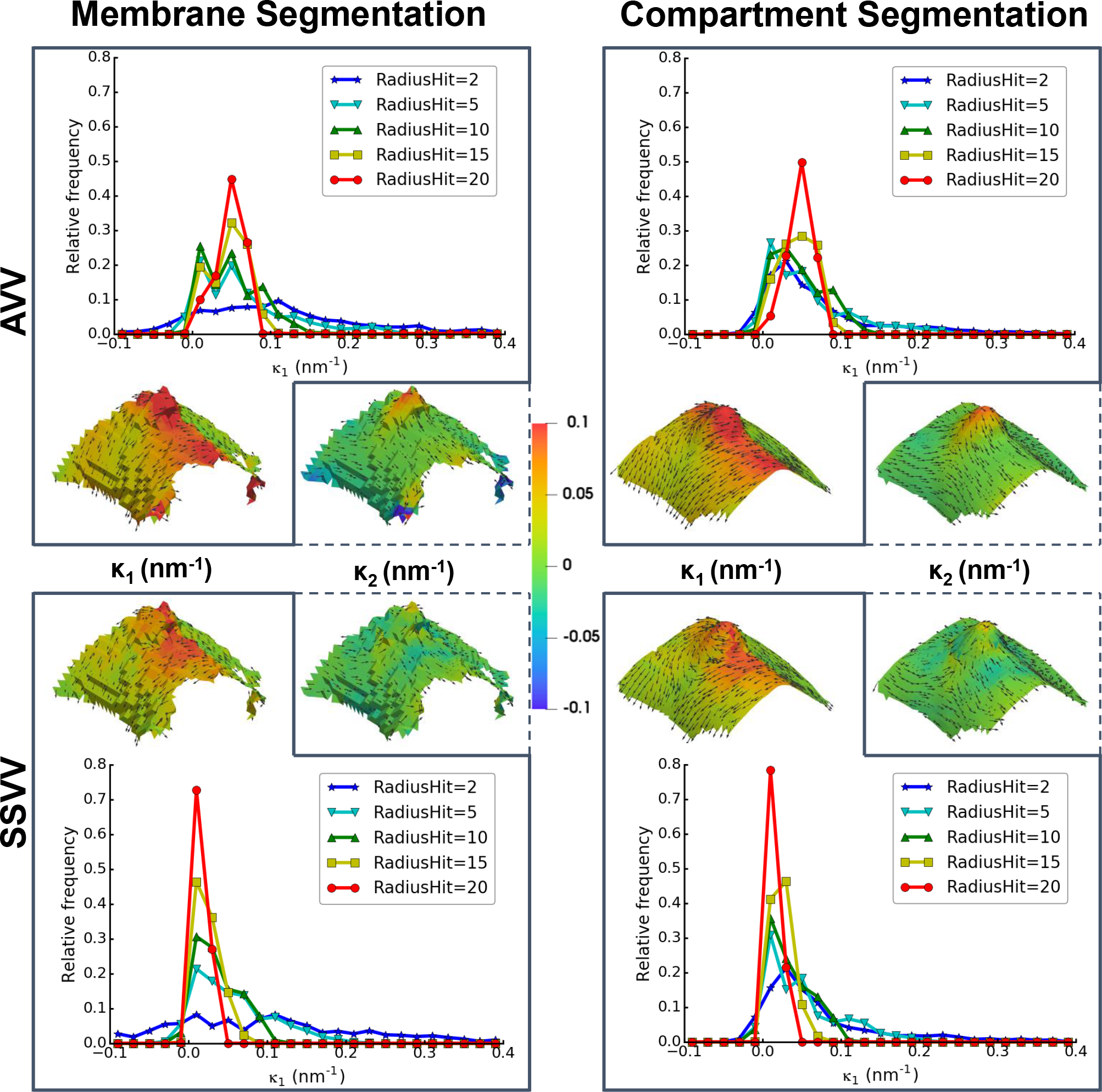
Methods and parameter comparison using a small cER membrane feature. Curvatures of a cER membrane region with base radius~10 nm were estimated on a surface generated using a membrane segmentation (left column) or a compartment segmentation (right column) by AVV (upper row) or SSVV (lower row) using *RadiusHit*=2, 5, 10, 15 and 20 nm. For the relative frequency histograms of estimated *κ*_1_ using the different *RadiusHit* parameters, values within 1 nm to surface borders were excluded. Visualizations of the estimated *κ*_1_ and *κ*_2_ (color scale was set to value range of [−0.1, 0.1]) are shown along with the corresponding principal directions (gray arrows, sampled for every forth triangle).

Then, we visualized the principal curvatures of the peak feature using *RadiusHit*=10 nm to analyze the difference between the two curvature estimation methods (visualizations in Fig. 9). We observe that real membrane features have a complex structure with a diverse curvature distribution with several local maxima and minima. For this specific feature, its principal curvatures estimated by AVV increase as we move from the base to the summit, while SSVV underestimates the curvatures, especially *κ*_2_. Since SSVV samples only surface points at distance=*RadiusHit* from each triangle center, the high curvature at and near the summit is overseen. Contrary to SSVV, AVV considers all triangles within the geodesic neighborhood and thus estimates the curvature increase towards the summit correctly. This example confirms that AVV performs better than SSVV for complex surfaces like biological membranes.

Moreover, we evaluated the effect of the quantization noise using the two surface types. The surface obtained directly from the membrane segmentation is step-like because of the quantization noise, which explains why curvature distributions for *RadiusHit*=2 nm are very broad (Fig. 9). In the surface based on compartment segmentation, the quantization noise has been to a great extend filtered, so even with *RadiusHit*=2 nm the distributions are similar to those using *RadiusHit*=5 and 10 nm.

#### 3.2.2 Border Filtering Decision

To conclude, we applied all optimal settings (surface extraction using the compartment segmentation, AVV and *RadiusHit*=10 nm), on the full cER surface (Fig. 10A), where the peak shown in Fig. 9 was cropped from. As already demonstrated using a cylinder surface in Sec. 3.1.7, our tensor voting-based methods make higher errors near surface borders. These errors can be especially high for AVV, since its range is not limited by *RadiusHit*. Therefore, we decided to exclude the borders of the membrane surfaces from the curvature calculation. At the same time, it is important to adjust precisely the distance from the graph border to minimize the percentage of discarded surface area. For the cER compartment segmentation with curvature estimated by AVV with *RadiusHit* = 10 nm (Fig. 10A), we made this decision using the diagram shown in Fig. 10B. It is necessary to filter out 5 nm from border in order to remove the extreme curvature values |0.4-0.8| nm^−1^;, while almost 80% of the surface still remains. The remaining curvature values are within the expected range of −0.1 to 0.15 nm^−1^; (Fig. 10C) and do not decrease much further when removing larger border regions (Fig. 10B).

**Figure 10:**
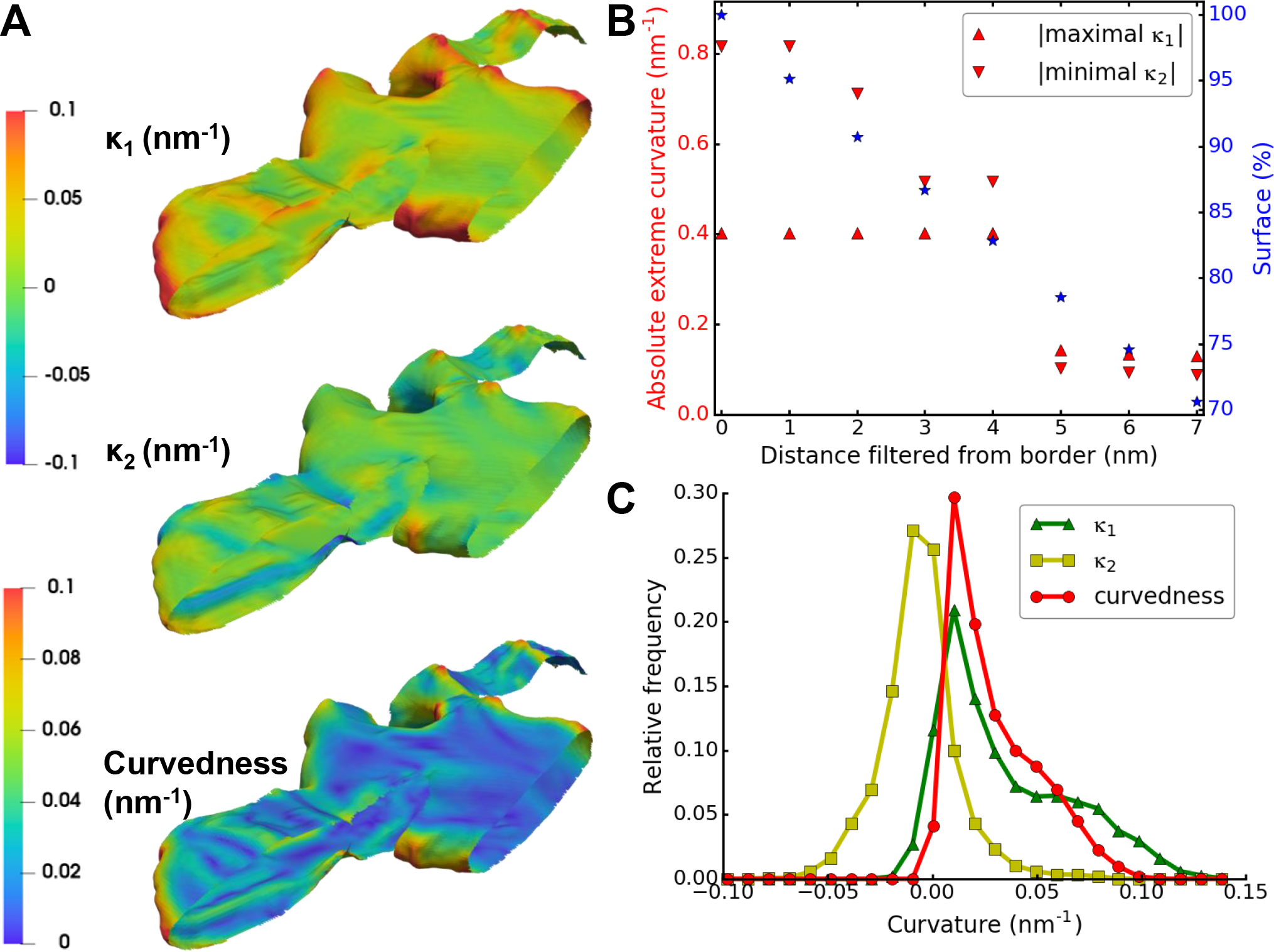
Application and border filtering decision for a cER membrane. Analysis of cER membrane curvatures, *κ*_1_, *κ*_2_ and curvedness, estimated by AVV with *RadiusHit* of 10 nm on the surface generated using the compartment segmentation. (**A**) Visualizations of the curvatures. Color scale was set to value range of [−0.1, 0.1] for *κ*_1_ and *κ*_2_ and of [0, 0.1] for curvedness. (**B**) Border filtering decision plot for the cER surface. For a distance filtered from border (X-axis), the absolute value of the curvature extremes (maximal *κ*_1_ and minimal *κ*_2_; red triangles) is shown on the left Y-axis, while percentage of the remaining surface area is shown on the right Y-axis (blue asterisks). (**C**) Relative frequency histograms of the curvatures. Values within 5 nm to surface borders were excluded and are not displayed in the histograms.

## 4 Conclusion

In this study, we described a procedure for local curvature measurement of biological membranes and validated it on artificial and cryo-ET data.

The first step is to represent the membranes as a triangle mesh surface that can be obtained from two different types of input: membrane segmentation or a filled segmentation of a membrane-bound cellular compartment. The second case usually demands more human intervention but orientation is recovered perfectly, and it contains less quantization noise because surfaces are generated at subvoxel precision. Surface triangles are mapped to a graph to facilitate the computation of geodesic distances.

The second step is to determine local curvatures and principal directions. Here, we tested the publicly available VTK [Schroeder et al., 2006], RVV (a modification of Normal Vector Voting from [Page et al., 2002] to estimate curvature sign correctly) and our two proposals AVV and SSVV (adaptations of [Page et al., 2002] and [Tong and Tang, 2005]). Tests using synthetic surfaces show that tensor voting-based approaches RVV, AVV and SSVV are more robust against quantization noise than VTK. AVV performs better than RVV for non-uniformly tessellated surfaces. For complex non-spherical surfaces, like biological membranes, AVV works better than SSVV.

Curvature is a local property, so its value on discrete surfaces depends on the definition of a neighborhood. Robustness increases with the size of the neighborhood by averaging the contributions of the contained triangles. However, features smaller than the size of the neighborhood are averaged out. Therefore, the neighborhood size defines the scale of the features that can be analyzed. To achieve more reliable results for cryo-ET segmentations that contain holes, curvature values at surface borders and/or higher than 1/*RadiusHit* should be excluded from the analysis.

We suggest that our method can be applied to any segmented membrane compartments or even other structures from which a surface can be extracted, not necessarily originated from cryo-ET data.

## Acknowledgment

We thank Markus Hohle for helpful discussions. M.K. and J.C. have been supported by the Graduate School of Quantitative Biosciences Munich. The authors have received funding from the European Commission (FP7 GA ERC-2012-SyG_318987–ToPAG).

